# DNA repair and mutagenesis in vertebrate mitochondria: evidence for the asymmetric DNA strand inheritance

**DOI:** 10.1101/842617

**Authors:** Bakhyt Matkarimov, Murat K. Saparbaev

## Abstract

A variety of endogenous and exogenous factors induce chemical and structural alterations to cellular DNA, as well as errors occurring throughout DNA synthesis. These DNA damages are cytotoxic, miscoding, or both, and are believed to be at the origin of cancer and other age related diseases. A human cell, in addition to nuclear DNA, contains thousands copies of mitochondrial DNA (mtDNA), a double-stranded, circular molecule of 16,569 bp. It was proposed that mtDNA is a critical target for reactive oxygen species (ROS), by-products of the oxidative phosphorylation (OXPHOS), generated in the organelle during aerobic respiration. Indeed, oxidative damage to mtDNA are more extensive and persistent as compared to that of nuclear DNA. Although, transversions are the hallmarks of mutations induced by ROS, paradoxically, the majority of mtDNA mutations that occurred during ageing and cancer are transitions. Furthermore, these mutations exhibit a striking strand orientation bias: T→C/G→A transitions preferentially occur on the Light strand, whereas C→T/A→G on the Heavy strand of mtDNA. Here, we propose that the majority of mtDNA progenies, created after multiple rounds of DNA replication, are derived from the Heavy strand only, due to asymmetric replication of the DNA strand anchored to inner membrane *via* D-loop structure.

## 1. Introduction

Mitochondria are organelles which exist in the form of an extensive tubular network and can be found in most of eukaryotic cells. These organelles have a complex mesh-like structure across the cytosol and undergo continuous cycles of fission and fusion that break and connect the network, respectively. This process enables the regulation of cellular signalling, homeostasis and redistribution of mitochondrial genetic materials. Mitochondria are cellular power plants which oxidize or “burn” carbohydrates, amino acids and fatty acids to synthesize adenosine triphosphate (ATP) in the process referred as the oxidative phosphorylation (OXPHOS) and electron transport chain (ETC). Mitochondria are double-membrane organelles composed of an outer membrane surrounding and an inner membrane which has five times bigger surface area than the outer one. The inner membrane is extensively folded into compartments referred as cristae. The space between the two membranes is termed the inter-membrane space and the space inside inner membrane is called matrix. The five main OXPHOS complexes are concentrated in the cristae membranes. The complexes I, III and IV pump proton ions H^+^ from the matrix to the inter-membrane space to generate a proton concentration gradient across the inner membrane. The complex V (ATP synthase) then uses this gradient to make ATP.

Mitochondria contain their own genomic DNA, referred as mitochondrial (mt) DNA with unique replication, transcription and translational machinery. Human cell contains several thousands copies of mtDNA, which is organized as a small closed circular duplex DNA of 16,569 bp. The human mitochondrial genome encodes 13 structural proteins which are essential subunits of the OXPHOS complexes I, III, IV and V; two mitochondrial ribosomal RNAs and 22 mitochondrion-specific tRNAs (Mechanic et al. 2006). Noteworthy, a mitochondrial genome employs a non-universal genetic code in which AUA is read as methionine, UGA is read as tryptophan, and AGA and AGG are read as STOP instead of arginines (Suzuki and Nagao 2011). An other intriguing particularity of the vertebrate mitochondrial genome is an unusual misbalance of nucleotides composition on the two strand of the mtDNA which leads to the separation into a Heavy (H) and a Light (L) strand upon ultracentrifugation in a cesium chloride gradient (Clayton 1996). It should be noted that H-DNA strand is G+T-rich and composed of 31% G, 13% C, 25% A and 31% T, whereas L-DNA strand is C+A-rich and composed of 31% C, 13% G, 31% A and 25% T.

The mtDNA is organized into nucleoids, a structural unit composed of DNA tightly packed with mitochondrial transcription factor A (TFAM) protein (Bogenhagen 2012). The majority of nucleoids consist of only a single mtDNA molecule (Kukat et al. 2015; Kukat et al. 2011). Nucleoids are attached to the inner membrane on the side of matrix and partitioned in distinct mitochondrial compartments cristae, to ensure that mtDNA are distributed evenly throughout the cell’s organelles (Kopek et al. 2012). Mitochondrial nucleoid is a platform for transcription and replication of mtDNA that ensures accurate replication and effective partition of the genetic materials. The replication of mtDNA is continuous and not dependent on the cell cycle which is essential for maintaining a high copy number of mtDNA per cell (Sasaki et al. 2017). It is commonly agreed that mtDNA is bound to the mitochondrial inner membrane by a mechanism involving the control region sequence in mtDNA (Jackson et al. 1996). After replication termination the separation of two newly synthesized molecules of mtDNAs is spatially linked to the division of the mitochondrial network, suggesting that mtDNA replication and mitochondrial fission are coupled processes (Garrido et al. 2003; Lewis et al. 2016). Fragmentation of the mitochondrial network *via* extensive fissions in G2 phase of cell cycle ensures the homogenous partition of nucleoids across the organelles before cell division and that daughter cells receive equal number of mtDNAs during mitosis (Mishra and Chan 2014). Thus the nucleoids in mitochondria of a cell are genetically uniform and they can segregate as a single unit during mitochondrial fission (Ban-Ishihara et al. 2013; Garrido et al. 2003; Lewis et al. 2016).

More than 90% of the oxygen used for cell respiration is consumed by the ETC in mitochondria. Premature electron leakage from the OXPHOS complexes I and III convert 1–2% of the oxygen to reactive oxygen species (ROS) namely superoxide (O2•^−^). The superoxide anions are converted to hydrogen peroxide (H_2_O_2_) by the superoxide dismutases. Non-detoxified H_2_O_2_ can be converted to the hydroxyl radical (HO•) *via* the Fenton reaction and then damage all cellular components (Murphy 2009). The nucleoids are anchored to inner membrane in close proximity to the ETC which is the main source of reactive oxygen species (ROS) within the cell. Although, the mtDNA packaging into nucleoide provides certain protection, it is believed that ROS would preferentially target mtDNA rather than nuclear genetic material. In the 1970s, Harman proposed that the ROS-mediated damage is an important contributor to somatic mtDNA mutations that result in production of dysfunctional ETC components, which in turn generate the increased level of ROS, thus leading to a “vicious cycle” responsible for aging (Harman 1972). This laid the base for the mitochondrial theory of aging which postulates that accumulation of the ROS-induced mtDNA mutations will lead to aging in humans and animals (Miquel et al. 1980). Studies in the past consistently showed more oxidative base damage in mtDNA as compared to that in nuclear DNA, in agreement with the hypothesis that endogenous damage accumulate preferentially in mtDNA (Ames et al. 1995; Hudson et al. 1998). Moreover, environmental factors predominantly target mtDNA than nuclear DNA, suggesting that the mitochondrial genome is more susceptible to damage induced by exogenous agents (Yakes and Van Houten 1997). Although oxidative damage accumulate in cellular DNA with age (Bokov et al. 2004), the evidence for causality of ROS-induced damage to mtDNA as a driving force behind aging is still lacking (Alexeyev 2009).

Early estimates of the mutation rate in mtDNA showed a 10-fold higher rate of evolution for mtDNA relative to the nuclear genome from somatic tissues of different primates studied (Brown et al. 1982), indicating a highly increased mutation rate for mtDNA. More recent estimates revealed 10- to 17-fold higher mutation rate of mtDNA than nuclear genome (Tuppen et al. 2010). It was suggested that the higher mutation rate in mtDNA is caused by ROS. Alternatively, other factors such as errors mediated by mitochondrial DNA replication machinery has been proposed as a primary source of mtDNA mutations (Zheng et al. 2006). MtDNA within a cell or individual can exist as a pure population of wild type (homoplasmy) or a mixture of wild type and mutant mtDNA, referred as heteroplasmy. Because of continuous cycles of fusion and fission that enable mixing and homogenisation of mitochondrial matrix, cells can maintain some level of heteroplasmy, in which functional and dysfunctional mtDNA variants coexist (Mishra and Chan 2014). For most of the known mtDNA mutations, no phenotypic alteration is ascribed unless mutant mtDNA molecules exceed 60% of the total mtDNA (Gilkerson et al. 2008; Schon and Gilkerson 2010). Nevertheless, the segregation of heteroplasmic population of mtDNA in cells could result in an unequal partitioning of mtDNA variants either at cell division, or by preferential replication of a specific mtDNA variant. Deep sequencing of mtDNA revealed the presence of low-level sequence variants in healthy individuals (Elliott et al. 2008; Payne et al. 2013), therefore if the proportion of mtDNA containing a dysfunctional mutation segregates above the threshold level, these heteroplasmic individuals might be at risk of developing a mitochondrial disease.

Recent advances in understanding the mechanisms of DNA replication, repair and mutagenesis of vertebrate mitochondrial genome revealed unexpected patterns that are incompatible with hypothesis of oxidative DNA damage driven mutagenesis. **In this review, we will discuss the basic mechanisms of mtDNA replication, repair and mutagenesis and then propose a hypothetical model which addresses some unresolved issues**.

## 2. Replication of mtDNA

Mitochondria depend on the nuclear genome for mtDNA replication and maintenance, since, all proteins required for replication, transcription, and repair of mitochondrial genome are encoded by the nuclear DNA (Bogenhagen 2012). Nevertheless, the DNA replication machinery of mitochondria in mammalians is different from that used for nuclear DNA replication and many mitochondrial replication factors are related to proteins identified in T3/T7 bacteriophages (Shutt and Gray 2006). DNA polymerase γ (PolG) is the high-fidelity replicative polymerase in mitochondria. Human PolG is a hetero-trimer consisting one 140 kDa catalytic subunit (PolG1) and two p55 accessory subunits (PolG2) (Fan et al. 2006; Gray and Wong 1992; Yakubovskaya et al. 2006). The catalytic subunit has 5’→3’ DNA polymerase, 3’→5’ proofreading exonuclease, and 5’-deoxyribosephosphate (dRP) lyase activities domains. The two accessory subunits are necessary for tight DNA binding to promote processive DNA synthesis (Young et al. 2011). TWINKLE is the mitochondrial DNA helicase which is required for strand unwinding and separation during mtDNA replication (Young and Copeland 2016). Mitochondrial Topoisomerase I (mtTop1) catalyzes transient cleavage and ligation of one strand of the duplex DNA in order to relieve tension and DNA supercoiling during replication (Korhonen et al. 2003; Korhonen et al. 2008; Spelbrink et al. 2001). Transcription is strongly coupled with DNA replication in mitochondria since mitochondrial RNA polymerase (POLRMT) can act as a primase in mtDNA replication (Fuste et al. 2010; Wanrooij et al. 2008), in addition to PrimPol, a primase-polymerase member of archaeo-eukaryotic primase superfamily (Rudd et al. 2014). Along with POLRMT, mitochondrial transcription factor A (TFAM) plays a central role in the mtDNA production of truncated transcripts from L-strand promoter (LSP) that are used to prime DNA synthesis during mtDNA replication (Shi et al. 2012). TFAM is a member of the high mobility group (HMG) box domain family and plays essential role in mtDNA packaging (Farge et al. 2014; Kukat et al. 2015). The single-stranded DNA (ssDNA)-binding protein (mtSSB) is an essential component of the mtDNA replisome. MtSSB binds to ssDNA and stimulates the activity of PolG (Farr et al. 1999; Korhonen et al. 2004). In addition to the above mentioned proteins, mitochondrial genome maintenance exonuclease 1 (MGME1) (Kornblum et al. 2013), DNA ligase IIIα (Lakshmipathy and Campbell 1999; Puebla-Osorio et al. 2006), Ribonuclease H1 (RNase H1) (Cerritelli et al. 2003; Holmes et al. 2015), DNA helicase/nuclease 2 (DNA2) (Copeland and Longley 2008), DNA flap-structure endonuclease 1 (FEN1) (Kalifa et al. 2009) and mitochondrial isoform of Topoisomerase IIIα (TopoIIIα) (Nicholls et al. 2018) are required for mtDNA maintenance and replication.

Three models for mtDNA replication have been proposed: the strand-displacement model (SDM) (Robberson and Clayton 1972), the ribonucleotide incorporation throughout the lagging strand model (RITOLS) (Yasukawa et al. 2006), and the strand-coupled model (Holt et al. 2000). In the commonly accepted SDM model, mtDNA replication occurs *via* asynchronous synthesis from distantly located sites O_H_ and O_L_. (Gustafsson et al. 2016). In the beginning, replication is initiated at O_H_ by PolG and a nascent H-strand is synthesized while the mtDNA is unwound by TWINKLE, and displaced parental H-strand is coated with mtSSB, preventing POLRMT-initiated transcription. After the H-strand (leading strand) replication fork passes O_L_, the later adapts a stem-loop structure, which prevents mtSSB binding, and instead is recognized by POLRMT. The RNA polymerase initiates primer synthesis and then is replaced by PolG for synthesis of the L-strand (lagging strand) (Fuste et al. 2010). After that the DNA synthesis continue on both H- and L-strand until the replication is terminated at O_H_ and O_L_, respectively. RNase H1 is essential for replication termination through primer removal. The last two ribonucleotides of the RNA-DNA junction, left after RNAse H1, are displaced by PolG into a flap-structure that is subsequently removed by DNA2, FEN1, and MGME1. The single-stranded break left after nucleases is sealed by DNA ligase IIIα. Intriguingly, the mtDNA replication termination results in the formation of a hemicatenane, two-ring interlocked DNA molecules at the O_H_ site *via* ssDNA linkage. Recently, it has been demonstrated that the decatenation of mtDNA after replication termination is catalyzed by TopoIIIα, which is essential for the segregation of nucleoids within the mitochondria (Nicholls et al. 2018). The mtDNA hemicatenane is unusual in the context of segregation of replicated circular double-stranded (ds)DNA molecules because this process in general proceeds *via* catenanes, DNA rings mechanically-interlocked *via* dsDNA linkage. At present, it is unclear how hemicatenane structures are generated during mtDNA replication, although the localisation of ssDNA linkage in hemicatenanes to the O_H_ region suggests that they are formed during the completion of mitochondrial genome replication.

In addition to protein components and specific regulatory DNA sequences, mtDNA contains unusual structures which participate in the regulation of DNA transcription and replication. For example, significant part of mtDNA molecules bear a third strand of DNA, referred as “7S DNA”, which generates a displacement (D) loop covering much of the control region (CR) referred also as major non-coding region (NCR) (Nicholls and Minczuk 2014). The D-loop spans approximately 600 bp between O_H_ and TAS regions of mammalian mtDNA and is present from around 10% of mtDNA in cultured human cells, up to 90% in Xenopus oocytes (Brown et al. 1978; Callen et al. 1983; Hallberg 1974). It is thought that the D-loop has many roles: it acts as a recruitment site for proteins involved in the organisation of mtDNA into nucleoid structures (He et al. 2007; Holt et al. 2007) and functions as a key component of replication (Antes et al. 2010; Clayton 1982). Recently, it has been demonstrated that D-loop has complex organization: it contains an RNA strand on the opposing strand to the 7S DNA forming an R-loop, which may have a role in the organisation and segregation of mtDNA (Akman et al. 2016). Intriguingly, cellular turnover of 7S DNA is very rapid with a half-life of around 1 h in rodent cells (Gensler et al. 2001). Depletion of MGME1 results in a large accumulation of 7S DNA suggesting that this ssDNA nuclease participates in the D-loop turnover (Kornblum et al. 2013; Szczesny et al. 2013).

## 3. DNA damage and repair of mtDNA

### 3.1. Nature of DNA damage

DNA damage can be classified by its nature: spontaneous *versus* induced; by structure: complex *versus* singular, bulky *versus* nonbulky, base *versus* sugar damage, single *versus* clustered damage; by biological consequences: innocuous *versus* toxic and/or mutagenic (Figure 1). Importantly, DNA can undergo spontaneous decomposition because of its intrinsic chemical instability. Spontaneous hydrolysis of DNA in water under physiological conditions results in purine loss and cytosine deamination at significant rates leading to appearance of abasic (AP) sites and uracil residues, respectively (Lindahl and Andersson 1972; Lindahl and Nyberg 1974). Under physiological condition 1/100,000 purines are lost from DNA every 24 hours resulting in abasic sites (Lindahl and Karlstrom 1973). Cytosine, adenine and guanine bases can undergo spontaneous loss of their exocyclic amino groups (deamination) resulting in highly mutagenic uracil, hypoxanthine and xanthine residues, respectively, that lead to C*→T, A*→G and G*→A transitions, respectively (Hill-Perkins et al. 1986; Kamiya et al. 1992). Under typical cellular conditions, deamination of DNA-cytosine to uracil occurs in about one of every 10^7^ cytidine residues in 24 hours, whereas guanine and adenine deamination occurs at 1/10 of this rate (Lindahl and Nyberg 1974). Importantly, deamination of C, A and G bases in single-stranded DNA occurs 10 times faster as compared to duplex DNA.

**Figure 1.**
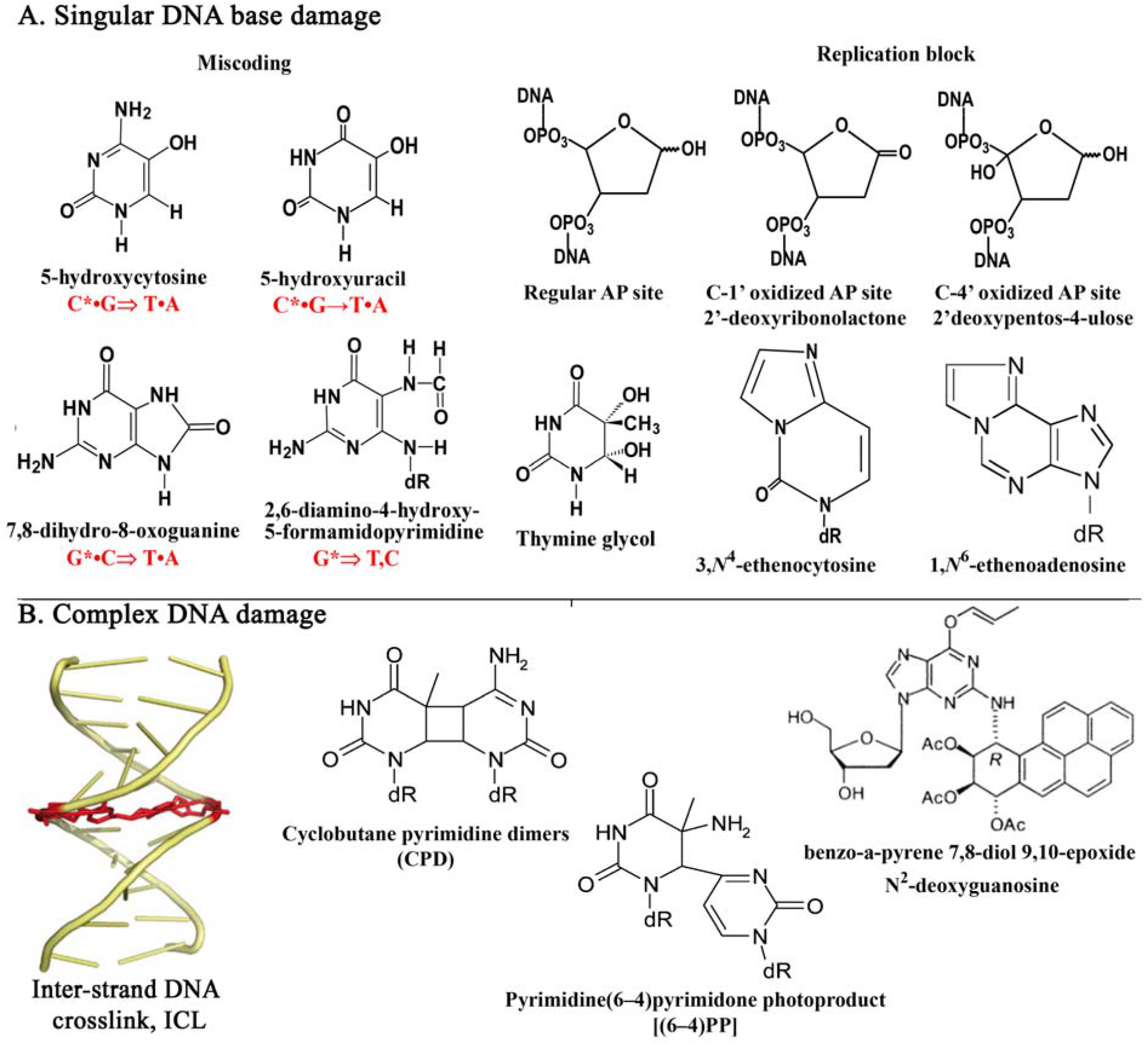
Schematic presentation of DNA damage induced by endogenous and exogenous factors. (**A**) Chemical structures of singular DNA base and sugar damages induced by oxygen free radicals. (**B**) Chemical structures of complex DNA damage having bulky and clustered characters.

ROS generated during aerobic respiration in mitochondria is a major source of endogenous DNA damage. Studies of oxidative damage to mtDNA revealed that oxidized bases occur more frequently, and persist longer in mtDNA, as compared to nuclear DNA damage (Richter et al. 1988; Yakes and Van Houten 1997). About 80 different types of base and sugar lesions induced by ROS have been identified (Bjelland and Seeberg 2003; Cadet et al. 2003) (Figure 1). ROS can damage nucleobases and sugar moieties in DNA either directly or indirectly. Hydroxyl radicals preferentially react with C8 atom of purines in DNA to generate 8-oxo-7,8-dihydroguanine (8oxoG), 8-oxo-7,8-dihydroadenine (8oxoA) and formamidopyrimidines (Fapy) (Cadet et al. 2003; Dizdaroglu 2012), and with C5-C6 double bond of pyrimidines to form glycols (Schuchmann et al. 1984; Téoule et al. 1977) (Figure 1). Abstraction of an hydrogen atom at the C1’ and C4’ positions of 2’-deoxyribose by ROS results in the formation of oxidized abasic sites: 2’-deoxyribonolactone and 2’-deoxypentos-4-ulose, respectively (Figure 1) (Dizdaroglu et al. 1977). The major endogenous oxidized bases 8oxoG, 5-hydroxyuracil (5ohU) and 5-hydroxycytosine (5ohC) are miscoding and, if not repaired, lead to mutation upon replication (Grollman and Moriya 1993; Kreutzer and Essigmann 1998; Kunkel and Bebenek 2000). C→T transitions and G→T transversions are the most common point mutation occurring in tumour suppressor genes commonly mutated in human cancers (Pfeifer 2000). C→T substitution could arise from mispairing of cytosine-derived lesions such as uracil, 5ohU and 5ohC with adenine (Kreutzer and Essigmann 1998). Whereas, G→T transversion results from mismatched pairing of 8oxoG present in the template DNA strand with adenine in newly synthesized strand (Grollman and Moriya 1993). Noteworthy, the steady state levels of 5ohU and 5ohC residues in DNA from mammalian tissues and human cells are higher than that observed with 8oxoG (Wagner et al. 1992). Oxidation of adenine residues in DNA results in the formation of 8oxoA and 2-hydroxyadenine (2-oxoA) (Kamiya et al. 1995). It should be noted that the formation of oxidatively induced adenine modifications including 8oxoA and FapyA is about 10-fold lower than that of related guanine degradation products upon exposure of cellular DNA to either hydroxyl radical or one-electron oxidants (Cadet et al. 2008; Cadet et al. 2010; Pang et al. 2014). Damage to the free nucleotide pool is also common and generates a similar spectrum of lesions (Cadet et al. 2003; von Sonntag 2006).

Indirectly, ROS can generate reactive aldehydes as products of membrane lipid peroxidation (LPO), which can react with DNA bases forming exocyclic etheno (ε) adducts 1,*N*^6^-ethenoadenine (εA) and 3,*N*^4^-ethenocytosine (εC) (Marnett and Burcham 1993) (Figure 1). Etheno adducts are ubiquitous and have been found in DNA isolated from tissues of untreated rodents and humans (Nair et al. 1995). Importantly, εA and εC levels are significantly increased by cancer risk factors related to oxidative stress/LPO, such as dietary ω-6 fatty acids intake, chronic infections and inflammatory conditions (Bartsch and Nair 2000). The εA and εC residues in DNA are highly mutagenic, εC mostly produces C•G→A•T transversions and C•G→T•A transitions (Basu et al. 1993; Moriya et al. 1994), whereas, εA residues are highly mutagenic in mammalian cells and result in T•A→A•T transversions (Levine et al. 2000; Pandya and Moriya 1996). Therefore the processes that prevent mutagenic effects of ε-adducts when present in DNA should play an essential role in maintaining the stability of mitochondrial and nuclear genomes.

Furthermore, cells are also exposed to environmental mutagens such as ionizing radiation, UV light, alkylation and DNA crosslinking agents. Ionizing radiations induce DNA strand breaks, oxidized AP sites and bases and clustered lesions. UV radiation generates two most common DNA lesions: the cyclobutane pyrimidine dimer (CPD) and the pyrimidine (6–4) pyrimidone photoproduct [(6–4)photoproduct; 6–4PP]. Both photoproducts are cytotoxic (block DNA replication and transcription) and mutagenic, while CPDs are several times more frequent than 6–4PPs (Douki et al. 2000) (Figure 1). A hallmark of UV mutagenesis is the high frequency of C→T transitions at dipyrimidine sites in DNA, possibly due to the extremely high deamination rate of cytosine residues within CPD sites in DNA (Peng and Shaw 1996). Mono-functional alkylating agents such as N-methyl-N’-nitro-N-nitrosoguanidine (MNNG) and methyl methanesulfonate (MMS) react with DNA bases to generate 7-methylguanine (7meG), 3-methyladenine (3meA) and *O*^6^-methylguanine (*O*^6^meG) which are the most abundant alkylation lesions (Pegg 1984; Singer 1976). The major adduct *O*^6^meG mispairs with thymine during DNA replication, resulting in G•C→A•T transitions (Swann 1990). Bifunctional alkylating agents can generate a covalent bond between nucleotides on opposite strands of a DNA duplex resulting in formation of interstrand DNA crosslink (ICL). The platinum compounds such as cis-diamminedichloroplatinum (II), also known as cisplatin, reacts with guanines and induces mainly DNA diadducts: 65% d(GpG) intrastrand cross-links, 25% d(ApG) intrastrand cross-links and 5–8% ICLs between the guanines in the sequence d(GpC). ICLs are highly lethal DNA lesions that block DNA replication, transcription and recombination by preventing strand separation (Figure 1).

### 3.2. DNA repair systems

DNA repair is essential for cell survival and maintenance of tissue homeostasis. Cellular organisms must constantly contend with endogenous and exogenous DNA damage and for this reason they have evolved multiple overlapping DNA repair systems to counteract the genotoxic effects of these insults. Modified base lesions and base mismatches are specifically recognized among vast majority of regular match bases by DNA glycosylases and apurinic/apyrimidinic (AP) endonucleases in the base excision repair (BER) and nucleotide incision repair (NIR) pathways, respectively (Couve-Privat et al. 2007; Krokan and Bjoras 2013) (Figure 2). In the BER pathway, a DNA glycosylase hydrolyses the *N*-glycosylic bond between the damaged base and sugar, leaving either an apurinic/apyrimidinic (AP) site or a single-stranded DNA break (Figure 2, steps 1 and 3). DNA glycosylases are classified as mono- and bi-functional based on their mechanism of action. Monofunctional DNA glycosylases cleave the *N*-glycosydic bond, releasing the modified base and generating an AP site (Cunningham 1997). The resulting AP sites are then incised at 5’ by major human AP endonuclease 1 (APE1/APEX1/HAP-1/Ref-1), which generates single strand break with 3’-OH and a 5’-blocking deoxyribosophosphate (dRP) group (Figure 2, step 4). Several studies showed that the full-length form of APE1 protein is present in the mitochondria of mammalian cells (Li et al. 2010; Tell et al. 2001; Vascotto et al. 2011). In fact, APE1 contains the mitochondrial targeting signal in the C-terminal part between residues 289–318 (Li et al. 2010). Bifunctional DNA glycosylases not only cleave the *N*-glycosydic bond, but also have an AP lyase activity that eliminates the 3’ phosphate (β-elimination) or 3’ and 5’ phosphates (β,δ-elimination) of the resulting AP site in a concerted manner. β-Elimination produces a nick flanked by a 3’-terminal α,β-unsaturated aldehyde and a 5’-terminal phosphate, whereas β,δ-elimination yields a single-nucleoside gap flanked by two phosphates (Cunningham 1997; Dodson et al. 1994). The 3’-terminal phosphoaldehyde and 3’-phosphate are then removed by APE1 (Chattopadhyay et al. 2006) (Figure 2, step 6) and PNKP (Tahbaz et al. 2012), respectively, allowing a DNA polymerase to fill the gap before DNA ligase IIIα seals the resulting DNA nick (Demple and Harrison 1994). Two mono-functional DNA glycosylases have been identified in mitochondria: uracil-DNA glycosylase 1 (UNG1), which excises uracil residues derived from cytosine deamination (Anderson and Friedberg 1980; Nilsen et al. 1997) and mismatch-specific adenine-DNA glycosylase (MUTYH), a homolog of the *E. coli* MutY DNA glycosylase which excises adenine incorporated opposite to 8oxoG in duplex DNA (Ohtsubo et al. 2000; Takao et al. 1999). In addition, two bi-functional DNA glycosylases have been characterized in mitochondria: 8oxoG-DNA glycosylase 1 (OGG1) which excises 8oxoG residues opposite to cytosine and FAPY residues in duplex DNA (de Souza-Pinto et al. 2001; Takao et al. 1998) and pyrimidine specific DNA glycosylase NTHL1, a homolog of *E. coli* endonuclease III, which excises oxidative pyrimidine lesions (Karahalil et al. 2003). Regarding the biological role of DNA glycosylases, animal DNA glycosylase knockout models *Ung1^−/−^*, *Ogg1^−/−^*, *Mutyh^−/−^*, *Nthl1^−/−^*, including also double mutants *Ogg1^−/−^ Mutyh^−/−^* and *Nthl1^−/−^ Neil1^−/−^* mice exhibit increased spontaneous mutation rate in nuclear DNA and carcinogenesis (Andersen et al. 2005; Chan et al. 2009; Nakabeppu et al. 2006; Xie et al. 2004). Paradoxically, the studies of knockout of *Ogg1^−/−^* (Itsara et al. 2014) and *Ogg1^−/−^ Mutyh^−/−^* double knockouts (Halsne et al. 2012) fail to observe an increase in the spontaneous mutation rate in mitochondrial genome. The intriguing question arises is then why the cell keep a specific repair system for mtDNA if it is not defending against induced or endogenous DNA damage (Pawar et al. 2018). Nevertheless, it has been shown that expression of the OGG1 protein protects cells form ROS (Dobson et al. 2002; Lia et al. 2018; Wang et al. 2011a). In addition, *Ogg1^−/−^* and *Neil1^−/−^* mice are prone to obesity and insulin resistance, possibly due to their specific role in the metabolic regulation *via* maintenance of mtDNA (Sampath et al. 2011; Sampath et al. 2012).

**Figure 2.**
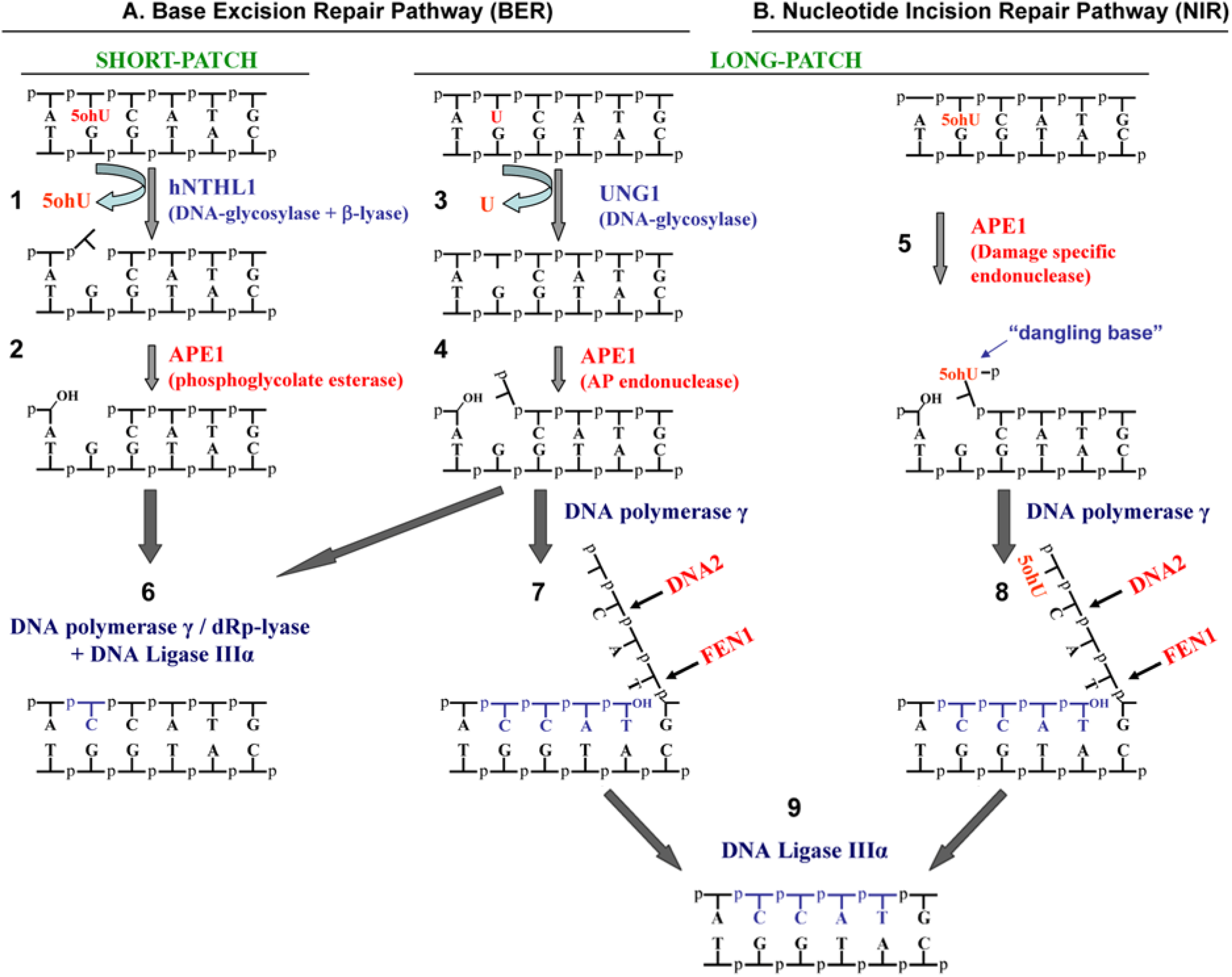
The base excision repair (BER) and nucleotide incision repair (NIR): two alternative DNA repair pathways for oxidative damage to mitochondrial DNA. (1-5) Upstream and (6-9) downstream steps of the BER and NIR pathways. In BER, (1) human bi-functional DNA glycosylase/AP lyase, hNTHL1 excises 5-hydroxyuracil (5ohU) residue in DNA resulting in formation of a free 5ohU base and single-strand break in form of one nucleotide gap containing a 3’-α,β-unsaturated aldehyde and a 5’-phosphate; (2) the 3’ repair diesterase activity of human APE1 removes 3’-blocking group to generate 3’-OH terminus; (2) human mitochondrial monofunctional uracil-DNA glycosylase 1, UNG1 excises uracil (U) residue in DNA resulting in formation of a free U base and abasic site (AP); (4) APE1 cleaves duplex DNA at the 5’ of AP site yielding a single-strand break with a 3’ hydroxyl adjacent to a 5’ deoxyribosephosphate (5dRp); In NIR, (5) APE1 directly cleaves 5’ next to 5ohU base generating single-strand nick containing a 3’-OH and a 5’-phosphate with dangling base. (6) In the SP-BER pathway DNA polymerase γ insert a single nucleotide and removes the 5’-dRp blocking residue, then DNA ligase IIIα seals the single-strand nick; (7-8) In the LP-BER and NIR pathways DNA polymerase γ initiates strand-displacement repair synthesis, coupled to DNA2- and FEN1-catalyzed cleavage of unannealed 5’-flap structures containing 5’-dRp residue in BER and 5’-dangling base in NIR pathway, respectively; (9) DNA ligase IIIα seals the single-strand nicks and restores genetic integrity of mtDNA.

In the downstream steps after sequential action of DNA glycosylases/AP endonucleases, two subcategories of the BER pathway are recognized: short- and long-patch BER (SP-BER and LP-BER, respectively) (Figure 2). In SP-BER only one nucleotide is removed upon damaged base excision, whereas in LP-BER more than one nucleotide is replaced with repair patch of 2-11 nucleotides. In the mitochondria DNA polymerase γ (PolG) removes blocking 5’-dRp residue by its AP lyase activity and then inserts single nucleotide (Longley et al. 1998) (Figure 2, step 6). However, dRp lyase activity of PolG is weak as compared to nuclear DNA polymerase β (Polβ), which may play a back up role in the mitochondrial SP-BER subpathway (Sykora et al. 2017). After filling of one nucleotide gap, the remaining nick is sealed by DNA ligase IIIα (LigIIIα) (Lakshmipathy and Campbell 2000). It should be noted, that the oxidized AP sites and cleavage products of the NIR pathway cannot be removed by the dRp-lyase function of PolG and Polβ, therefore these lesions are removed in LP-BER (Figure 2 steps 7-8). Noteworthy, among oxidative DNA damage generated upon oxidative stress, the oxidized AP sites might represent majority of sugar damage (Roginskaya et al. 2005) suggesting that SP-BER might be cytotoxic (Demple and DeMott 2002). During LP-BER, PolG initiates strand-displacement synthesis and generate 6-9 nucleotides ssDNA flap (Szczesny et al. 2008) containing the 5’-terminal blocking groups such as oxidized dRp residue or nucleobase, which is removed by the concerted action of two nucleases, FEN1 and DNA2 (Duxin et al. 2009; Kalifa et al. 2009; Zheng et al. 2008) (Figure 2 steps 7-8). The resulting nick is sealed by LigIIIα (Figure 2 step 9). Intriguingly, the LP-BER subpathway resembles to Okazaki fragment maturation and employs replication associated nucleases. Given the occurence of oxidized AP sites and strong strand displacement activity of PolG, the LP-BER subpathway might be the predominant mode of repairing oxidative DNA base damage in mitochondria.

The DNA glycosylase-initiated BER pathway raises problems since it generates genotoxic abasic and 3’-blocking group intermediates. Findings that AP endonucleases can directly cleave DNA 5’ to various oxidatively damaged nucleotides, generating 3’-hydroxyl and 5’-phosphate termini, together with genetic data on the cell resistance to oxidative stress, are suggestive of the existence of an alternative to the classic BER, referred as the NIR pathway, which bypasses the abasic intermediates (Ischenko and Saparbaev 2002) (Figure 2 step 5). AP endonucleases contain multiple repair activities and participate in the both BER and NIR pathways. AP site cleavage (or AP endonuclease) and 3’-repair phosphodiesterase activities can be referred as the BER functions, whereas nucleotide incision activity as the NIR function of the AP endonucleases (Gros et al. 2004). Under low concentrations of Mg^2+^ (≤1 mM), APE1 switches its substrate specificity and recognize diverse types of DNA base lesions including α-anomeric nucleotides (dN), oxidized pyrimidines such as 5,6-dihydrouracil, 5ohU and 5ohC (Daviet et al. 2007; Gros et al. 2004) (Figure 2 step 5), formamidopyrimidines (Christov et al. 2010), exocyclic DNA bases, thymine glycol, uracil (Prorok et al. 2013; Prorok et al. 2012), and bulky lesions such as benzene-derived DNA adducts (Guliaev et al. 2004). Interestingly, the study of subcellular localisation of APE1 showed varying patterns from mainly cytoplasmic to mixed cytoplasmic/nuclear and to mainly nuclear localisation (Kakolyris et al. 1998). Intriguingly, in the cell types with high metabolic or proliferative rates, APE1 is predominantly localized in the mitochondria and endoplasmic reticulum (Tell et al. 2001; Tell et al. 2005; Tomkinson et al. 1988). Furthermore, when cells are exposed to oxidative stress conditions, the level of APE1 in mitochondria is significantly increased in a dose- and time-dependent fashion (Frossi et al. 2002; Mitra et al. 2007), implying that APE1 participates in the maintenance of mtDNA under conditions of high energy metabolism and oxidative stress. In agreement with this, it has been demonstrated that down-regulation of the APE1 expression in mouse embryonic fibroblasts (MEF) resulted in the reversible suppression of the mitochondrial respiration and OXPHOS activity (Suganya et al. 2015). Based on these observations, we may speculate that in mitochondria APE1 initiates the DNA glycosylase-independent NIR pathway and shifts the removal of oxidized bases and AP sites into long patch repair synthesis similar to that in the LP-BER subpathway (Figure 2 steps 7-9).

Bulky DNA adducts, DNA-DNA and DNA-protein crosslinks are substrates for nucleotide excision repair (NER) pathway. In the NER pathway, a multiprotein complex recognizes and excises bulky DNA adducts in the form of short oligonucleotides that contain the lesion (Marteijn et al. 2014). NER is a major repair system that removes DNA damage induced by UV, anticancer agents such as cisplatin, and many other environmental carcinogens. In eukaryotic cells, NER involves dual incisions that bracket the lesion site and release 24-32 nucleotide-long oligomer containing the damaged residues. At present, the majority of *in vitro* and *in vivo* evidences indicate that CPD and 6-4 PPthymidine dimers (Clayton et al. 1975), cisplatin intrastrand crosslinks, complex alkylation damage, and other forms of damage (LeDoux et al. 1992; Pascucci et al. 1997), are not repaired in mtDNA implying the absence of NER in the mitochondria.

Base mispairs and short deletion-insertion loops are generated during DNA replication and homologous recombination. Mismatch repair (MMR) is an evolutionarily conserved system that recognizes and repairs mismatches in a strand-specific manner. MMR machinery is coupled to DNA replication and can distinguish the newly synthesized strand from the parental strand. In human cells two major heterodimers: Msh2/Msh6 (MutSα) and Msh2/Msh3 (MutSβ) recognize DNA mismatches and trigger their removal by recruiting MutLα (MLH1/PMS2) and MutLβ (MLH1/PMS1) complexes (Jiricny 2006). At present, the main proteins involved in nuclear MMR have not been found in mitochondria. Nevertheless, the repair factor YB-1, which contains a binding activity towards mismatched DNA, has been identified in mitochondria, and its knockdown decreases the MMR activity in the organelle (de Souza-Pinto et al. 2009).

### 3.3. Alternative pathways of mtDNA maintenance

For the reason that thousands copies of mtDNA are present in a cell, a significant loss of damaged DNA can be compensated by replication of remaining non-damaged molecules. Several studies have demonstrated that ROS induce the degradation of mtDNA and mtRNA (Abramova et al. 2000; Furda et al. 2012; Rothfuss et al. 2009; Shokolenko et al. 2009). Noteworthy, low levels of 8oxoG residues were detected in the circular mtDNA, whereas linear fragmented mtDNA contains very high 8oxoG, which is further increased after oxidative stress (Suter and Richter 1999). Therefore, the absence of significant mtDNA mutagenesis under oxidative stress conditions suggests that in mitochondria most of damaged DNA molecules are degraded perhaps due to the lesion-dependent transcription and replication blockage (Shokolenko et al. 2009). Noteworthy, it was shown that mtDNA, but not nuclear DNA, is also resistant to mutagenesis induced by powerful carcinogens, such as MNNG and ethylmethane sulfonate, most likely due to extensive degradation of the alkylated mtDNA (Marcelino et al. 1998; Mita et al. 1988). Based on these observations it has been proposed that the lesion-dependent DNA degradation can be considered as a specific mitochondrial repair pathway (Alexeyev et al. 2013). Indeed, the ectopic expression of certain isoforms of OGG1 protects cells against increased oxidative stress (Dobson et al. 2002; Lia et al. 2018), maintain normal neuronal biogenesis (Wang et al. 2010) and promote mitochondrial biogenesis during cell differentiation (Wang et al. 2011b). Thus, we may hypothesize that the DNA glycosylases-intiated BER pathway helps to destroy damaged mtDNA molecules, rather than repair them. This may explain the absence of somatic enrichment for transversion mutations in mtDNA and the role of DNA glycosylases in metabolic regulation in mice (Sampath et al. 2011; Sampath et al. 2012).

Finally, it should be noted that the tight packaging of mtDNA into nucleoid structure might offer an efficient protection against exogenous DNA damaging agents (Alan et al. 2016). The degree of compaction of mtDNA with TFAM from fully compacted nucleoids to naked DNA regulates transcription, replication and possibly also DNA repair and degradation *via* regulated access to DNA (Farge et al. 2014).

## 4. Mutagenesis of mtDNA

In animal models and humans point mutations and large deletions in mtDNA increase in frequency with age and have been implicated in the origin of age-related diseases (Greaves et al. 2014; Hahn and Zuryn 2019; Li et al. 2015; Sun et al. 2016). It is generally agreed that in a cell: mutations accumulates faster in mtDNA, as compared to that in nuclear DNA. The high replicational turnover of mtDNA is likely a main contributor to the increased spontaneous mutation rate because of the inevitable introduction of DNA polymerase errors during synthesis (Kennedy et al. 2013; Radzvilavicius et al. 2016). Importantly, the types of mtDNA mutations are cell type and age specific. Large deletions are more likely to accumulate in non-dividing cells: muscle fibers and neurons, but not in actively proliferating cells, such as colon mucosa and cancer cells (Khrapko and Turnbull 2014). Interestingly, pigmented neurons of substantia nigra in the brain of old individuals contain very high proportion of mtDNA deletions, but mtDNA from other types of neurons from the same subject lack deletions (Kraytsberg et al. 2006). Recent breakthroughs in DNA sequencing technology allowed detection of rare/sub-clonal mutations on genome wide level and characterize mutation spectra in mtDNA during aging and cancer (Ju et al. 2014; Kennedy et al. 2013; Williams et al. 2013). Study of the somatic mutations in mtDNA isolated from the pre-frontal cortex of human brain from young and old individuals revealed 5-fold increase in the frequency of point mutations in 80 years old group (Kennedy et al. 2013). Majority of point mutations in both the young and old samples were transitions, whereas transversions, mutational signature of oxidative DNA damage, were minor. Intriguingly, mutations accumulated asymmetrically on the two Heavy and Light strands of mtDNA. In young human brain samples C→T mutations were more likely to occur in H-strand (Kennedy et al. 2013). In the brains of aged individuals, this pattern became more prevailing, and was accompanied by T_L_→C_L_ mutations in the L-strand, mirroring the nucleotide composition bias of the light strand for A (~ 31%) over T (~ 25%). Quite unexpectedly, this strand asymmetry bias was reversed in the mtDNA’s control region which include H-strand origin of replication (O_H_), promoters for transcriptions and D-loop. This asymmetrical pattern of mtDNA mutation in the human brain has been confirmed in a subsequent DNA sequencing study (Williams et al. 2013). Furthermore, previous study of mtDNA mutations in human lung epithelium also showed that G and T sites in L-strand were underwent transition mutations more frequently than C and A sites (Zheng et al. 2006).

Previously, it was proposed that somatic mutations in mtDNA may contribute to tumour development by fulfilling increased energy demand of the uncontrolled cell proliferation associated with cancer (Wallace 2012). Recently, to address the role of mitochondrial mutations in cancer, Ju and colleagues have examined mtDNA sequences from 1675 cancer biopsies across 31 tumour types and compared to normal tissue from the same patients (Ju et al. 2014). In total 1907 somatic point mutations were identified, which exhibited similar replicative strand bias described above, with the prevalence of C_H_→T_H_ and A_H_→G_H_ transitions on the H-strand and T_L_→C_L_ and G_L_→A_L_ substitutions on the L-strand. Importantly, this mitochondrial pattern of somatic mutations differs from those identified in nuclear genome and most likely do not provide selective advantage to cancer cells. Furthermore, ROS, cigarette smoke and UV light had no or very little effect on mtDNA mutations in cancer, suggesting that an endogenous mutational mechanism linked to mtDNA replication is the major cause of mutagenesis in mitochondria (Ju et al. 2014). In their model Ju and colleagues proposed that during asynchronous replication of mtDNA, displaced, single-stranded H-strand is prone to cytosine and adenine deamination thus generating C_H_→T_H_/A_H_→G_H_ substitutions. Whereas, on the L-strand template PolG predominantly generates T_L_•G and G_L_•T mismatches which result in T_L_→C_L_/G_L_→A_L_ transitions (Ju et al. 2014). Remarkably, this strand bias is reversed in the D-loop region of mtDNA (Ju et al. 2014) and similar reversion in the mutation pattern occurs in mtDNA of aging brain described above (Kennedy et al. 2013). This observation point to the important feature of mtDNA replication, since inversion of Control Region in mtDNA of certain species of fish lead to a reversal of the strand bias in mutation pattern (Fonseca et al. 2014; Fonseca et al. 2008). Importantly, the change in strand-specific compositional bias in the mitogenomes containing the inverted Control Region is in agreement with asynchronous mode of mtDNA replication (SDM and RITOLS), but not with the Stand Coupled Model (SDM).

## 5. A hypothesis on the origin of mtDNA mutations

Here we propose a model of the asymmetric DNA strand inheritance, which provides a simple explanation to the observed highly biased pattern of mutations in vertebrate mtDNA (Figure 3). Our model is consistent with the strand-asynchronous models of mtDNA replication: SDM and RITOLS (Falkenberg 2018; Holt and Reyes 2012; Pohjoismaki et al. 2018), but contain several new, important features which apparently have not been discussed before. In this model we suggest that the single-stranded H-strand, anchored to inner membrane through specific interactions with Control Region or probably D-loop structure, would replicate in uninterrupted manner to make new DNA progeny (containing parental H-strand and nascent L-strand) (Figure 3, step 6). While, the L-strand will undergo only one cycle of replication which will make a new mtDNA molecule (containing parental L-strand and nascent H-strand) not attached to inner membrane (Figure 3, step 5). Thus, the L-strand derived progeny will not proceed to the next DNA replication cycle and would probably diffuse in mitochondrial matrix as non-attached nucleoids. In our highly asymmetric DNA strand replication model, the H-strand would produce majority of mtDNA offspring and the L-strand would make only minor fraction of mtDNA progeny. Nevertheless, we hypothesize that the pool of free and non-membrane-bound mtDNA progeny generated from L-strand replication may compete for a specific membrane attachment site required for mtDNA replication. In the SDM and RITOLS models of mtDNA replication: the synthesis of nascent L-strand on H-strand template from O_L_ is delayed compared with the synthesis of nascent H-strand on L-strand template from O_H_ (Figure 3, steps 2-3). Here, we suggest that the replication of H-strand is terminated with the DNA nick sealing before the completion of L-strand replication and this should generates hemicatenanes (two intertwined circular DNA associated through a ss linkage) (Figure 3, step 4), rather than catenanes (two intertwined circular DNA associated through a ds linkage). In agreement with our hypothetical model, recently it has been demonstrated that the mtDNA replication termination occurs *via* a hemicatenane formed at the O_H_ site and that TopoIIIα is essential for resolving this structure (Nicholls et al. 2018). It should be noted that in the past, Laurie and colleagues proposed a similar mechanism of the generation of hemicatenanes through alternative replication termination of circular DNA molecules (Laurie et al. 1998). In their model authors proposed that when two replication forks converge on the circular DNA, one of the advancing DNA polymerases becomes displaced from the template by the accumulated stress, so that only one fork advances. This induces a branch migration in which both the leading and lagging ends of DNA strands of the abandoned fork are progressively displaced. When the displaced DNA strand ends anneal back to complementary single-stranded regions of the advancing fork, the replication of both strands can be finished, and this results in hemicatenation (Laurie et al. 1998). Furthermore, formation of plasmid DNA hemicatenans in *Xenopus* egg extracts can be triggered by the DNA polymerase inhibitor aphidicolin which most likely provoke the asynchronous mode of DNA replication (Lucas and Hyrien 2000).

**Figure 3.**
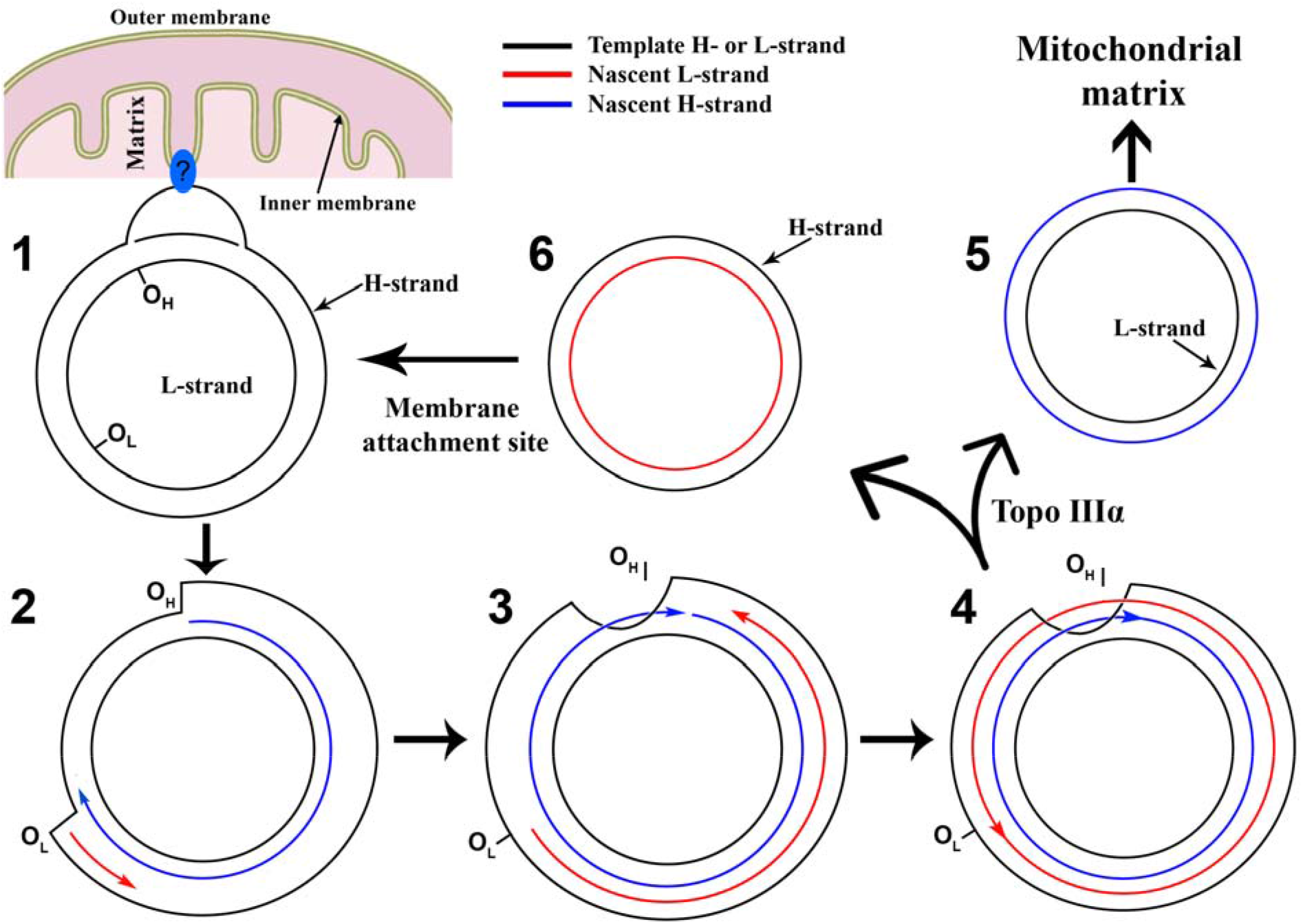
The model of mtDNA replication leading to asymmetrical DNA strand inheritance. (**1**) The mtDNA is anchored to inner membrane of mitochondria *via* interaction of D-loop region with a specific site in the membrane; (**2**) The asynchronous replication of leading H-strand (blue arrow) at O_H_ site proceeds unidirectionally to displace parental H-strand. When the O_L_ site is exposed, the lagging L-strand (red arrow) replication initiates in opposite direction; (**3**) Replication of leading H-strand proceeds further and terminates at O_H_ before the lagging L-strand reach this site. Premature termination of H-strand synthesis produces a hemicatenane: two-ring molecules mechanically interlocked *via* single-stranded linkage (Laurie et al. 1998); (**4**) Replication of lagging L-strand terminates at O_L_ site to complete the formation of a double-stranded hemicatenane composed of two interlocked mtDNA molecules. The hemicatenane is unlinked by mitochondrial Toposiomerase IIIα to produce two separate mtDNA molecule one containing parental H-strand and the other parental L-strand; (**5**) The mtDNA molecule containing old parental L-strand diffuses freely across mitochondrial matrix and may anchor to inner membrane through the expression of OXPHOS complex proteins (Lynch and Wang 1993); (**6**) The mtDNA molecule containing old parental H-strand stays attached to a specific site in the inner membrane and continues to replicate.

Our model also provides a simple explanation for the unusual strand bias in the somatic mutations observed in mitochondrial genome in aging brain and cancer (Ju et al. 2014; Kennedy et al. 2013). According to our model the prevalence of C→T/A→G transitions on the H-strand and absence or rare appearance of these type of substitutions (C_L_→T_L_/A_L_→G_L_) on the L-strand is due to the fact that majority of mtDNA progeny in vertebrate are derived from the H-strand, and that the propagation of mutations occurring in the L-strand is very limited. We will refer to this phenomenon as an asymmetric DNA strand inheritance. We suggest that a spontaneous cytosine deamination occurs on both mtDNA strands with similar rates and following of a DNA polymerase insertion of A opposite to U this should induce C→T transitions in both H- and L-strand. However, the mutation occurred in H-strand, but not that in L-strand, will be spread in the progeny due to the asymmetric DNA strand inheritance. This is why we observe the prevalence of C→T transitions on the H-strand and their mirror counterparts G→A transitions on the L-strand. Furthermore, it has been suggested that A→G transitions in mtDNA are due to the increased rate of deamination of adenines in the H-strand when it is exposed in single-stranded form (Ju et al. 2014). However, spontaneous deamination of both A and G occurs at 50-fold slower rate than that of C (Lindahl 1979), whereas *in vivo* A_H_→G_H_ transitions occurs at the rates comparable to that of C_H_→T_H_ (Ju et al. 2014; Kennedy et al. 2013). Also, deamination of G forms xanthine residue which can mispair with T, however, G_H_→A_H_ transitions occurs at much lower frequency than A_H_→G_H_. It should be stressed, that in addition to cytosine deamination, the most frequent spontaneous lesion in DNA is an abasic site (AP) which results from depurination. The rate of spontaneous formation of AP sites is at least tenfold greater than that of deamination of cytosines (Lindahl 1993). It is tempting to speculate that majority of A_H_→G_H_ transitions in mtDNA are due to spontaneous formation of AP sites. Noteworthy, yeast and human REV1, a translesion synthesis (TLS) DNA polymerase, preferentially inserts C opposite AP sites (Choi et al. 2010; Haracska et al. 2002) and that in budding yeast, dCMP is inserted (“C-rule”) opposite the AP-site in the single-stranded gap of a duplex plasmid (Gibbs and Lawrence 1995). According to “C-rule” the loss of adenines in mtDNA would give rise A→G transitions, while the loss of guanines would be non-mutagenic (Zhang et al. 2006). However, recent study demonstrated that enzymatically created AP sites in mtDNA of mouse cells are weakly mutagenic, and that repair and DNA degradation occur more often than TLS of AP sites (Kozhukhar et al. 2016). More studies are required to identify a possible origin of A_H_→G_H_ transitions in mtDNA.

The completion of mitochondrial replication cycle takes approximately 1 hours (Berk and Clayton 1974), indicating that the speed of DNA synthesis is much slower than that of eukaryotic nuclear and bacterial genomes (Clayton 1982). The time-consuming and asynchronous mode of mtDNA replication may ensure a high-fidelity copying of DNA, while the synchronous strand-coupled mechanism (SDM) is more fast and error-prone, but enables quick restoration of copy number after DNA damage (Torregrosa-Munumer et al. 2015; Yasukawa et al. 2005). In the RITOLS model: the replication of nascent H-strand starts early and generates partially replicated mtDNA in which displaced parental H-strand is covered by RNA. It should be noted that H-strand is the template from which most mitochondrial proteins (12 out of 13) are transcribed, whereas only one protein-coding gene, ND6, is transcribed from the L-strand. One may suggest that the H-strand is preferentially anchored to inner membrane *via* expression of the membrane-bound mitochondrial OXPHOS proteins (Lynch and Wang 1993) (Figure 3, step 1). Also, high level of transcription may inhibit mitochondrial replication, since the mtDNA deletion mutants acquire selective advantage in aging tissues (Kowald and Kirkwood 2014). Furthermore, we may speculate that the intensive transcription of H-strand serves to screen this portion of mtDNA for damage. If DNA transcription on the H-strand is blocked by a lesion then this may inhibit replication start from O_H_ site *via* the interference with D-loop structure. This mechanism of transcriptional DNA scanning may prevent the replication machinery from having to copy damaged DNA templates and thus keeping away from mutation fixations.

In summary, our model of the asymmetric DNA strand replication of vertebrate mtDNA explains the formation of hemicatenanes and unusual mutation pattern. Also, the proposed model may provide some insights into a putative evolutionary origin of cellular organelles. The ssDNA bacteriophages such as M13 and φX174 share similarities with vertebrate mtDNA in the asymmetric mode of DNA replication and strand nucleotide compositional bias, suggesting that the asymmetric strand replication may be a cause of the bias in strand nucleotides composition. Furthermore, the key proteins of the mitochondrial transcription and replication machineries are derived from the bacteriophages such as T3/T7 lineage of coliphages (Filee and Forterre 2005; Filee et al. 2002), some cyanophages (Chan et al. 2011) and others phages rather than from an α-Proteobacterium as thought before (Shutt and Gray 2006). Based on these observations, we may speculate that mitochondria and some bacteriophages are remnants of the extinct ancient free-living cellular organism which had a different type of genetic code. Mutational pressure and natural selection during billion years evolution lead to the emergence of a single universal genetic code in almost all free-living organisms and to the extinction of all other species of cellular organisms with alternative codes.

## 6. Conclusions

In this chapter we have reviewed the roles of DNA damage, DNA repair and DNA replication in the maintenance of mitochondrial genome in vertebrates. To summarize our review, we would like to highlight several essential points:

- Replication of mtDNA proceeds through asymmetrical replication of H-strand resulting in the predominant, biased inheritance of one strand in mtDNA;
- Asymmetrical DNA strand replication leads to an asymmetry in the somatic mtDNA mutational signature which in turn might be an evolutionary cause of the strand nucleotides compositional bias of animal mitochondrial genome;
- Formation of hemicatenanes during mtDNA replication suggest the premature termination of nascent H-strand replication;
- Studies of DNA damage, DNA repair and mutagenesis in mitochondria may suggest that the spontaneous decay of mtDNA is a major source of endogenous mutations rather than a DNA polymerase’s errors;
- A non-universal genetic code of animal mitochondria and similarities of mtDNA transcription and replication machineries with that of bacteriophages may point to the extinct ancient cellular organism with alternative DNA coding system as a putative mitochondrial ancestor.

## Acknowledgements

This work was supported by grants to Murat Saparbaev from la Ligue National Contre le Cancer “Equipe Labellisee”, Electricité de France (RB 2017) and French National Center for Scientific Research (PRC CNRS/RFBR n1074 REDOBER); and to Bakhyt T. Matkarimov from Nazarbayev University ORAU grant and MES RK grants АР05133910, AP05134683 program BR05236508.

